# hnRNPA1/2 homolog *hrpa-1* coordinates with miRNAs to regulate gene expression during *C. elegans* development

**DOI:** 10.1101/2022.07.22.501200

**Authors:** Shilpa Hebbar, Ganesh Panzade, Anna Zinovyeva

## Abstract

microRNAs (miRNAs) are small non-coding RNAs that play crucial roles in development and in disease. miRNAs associate with Argonaute proteins to form miRNA Induced Silencing Complexes (miRISCs), which post-transcriptionally repress gene expression. miRNA-mediated gene repression itself is subject to regulation by factors that can affect miRNA biogenesis or function. We previously identified HRPA-1, an hnRNPA/B homolog, as a putative physical interactor of miRNAs. Here, we report characterizations of both physical and genetic interactions between HRPA-1 and miRISC components. We confirmed HRPA-1 precipitation in let-7 and miR-58 pulldowns and detected an interaction between HRPA-1 and Argonaute. Deletion of *hrpa-1* in a *mir-48 mir-241(nDf51)* background enhanced the *mir-48 mir-241* developmental defects, suggesting that *hrpa-1* may be important for *let-7* family miRNA activity. Similarly, loss of *hrpa-1* strongly enhanced developmental defects associated with two other miRNA mutants, *lsy-6(ot150)* and *let-7(n2853)*. Depletion of HRPA-1 modestly disrupted miRNA levels and affected global gene expression profiles. We identified a potential target of *hrpa-1, R06C1*.*4*, whose knockdown partially recapitulates the *hrpa-1(-)* effects on miRNA mutant phenotypes. Overall, we demonstrate *hrpa-1* and *R06C1*.*4* roles in *C. elegans* developmental timing regulation and propose models describing possible coordinating modes of gene regulation by HRPA-1 and miRNAs.

## Introduction

Gene regulatory mechanisms orchestrate developmental and physiological processes by controlling spatiotemporal gene expression programs. microRNAs (miRNAs) and RNA binding proteins (RBPs) are essential components of the gene regulatory machinery, with both classes of molecules post-transcriptionally controlling gene expression. miRNAs bind target mRNAs in the 3’ UTR region via partial complementary base pairing, placing the miRNA induced silencing complex (miRISC) on the target RNAs to initiate translational inhibition and mRNA destabilization. RBPs exert their gene-regulatory influence through regulation of the RNA lifecycle. Coordination between miRNAs and RBPs contributes to the gene regulatory outcomes of gene expression programs. Heterogenous nuclear ribonucleoproteins (hnRNPs) are a family of closely related RNA binding proteins that associate with different classes of RNAs (Dreyfuss *et al*. 1993). HnRNPs harbor modular RNA binding domains and Proline or Glycine-rich regions (Dreyfuss *et al*. 2002). Members of this RBP family have been reported to affect alternative splicing, mRNA localization and mRNA stability among other processes (as reviewed in Geuens *et al*. 2016). A few prominent members of the family such as hnRNPA1, and hnRNPA2B1 have also been shown to bind DNA within the telomeric ends to regulate their stability (Jean-Philippe *et al*. 2013).

hnRNPs have also been implicated in miRNA-mediated gene regulation (Li *et al*. 2019; Shin *et al*. 2017; Fan *et al*. 2015; Guil and Ca’ceres, 2007; Michlewski and Ca’ceres, 2010; Kooshapaur *et al*. 2018). hnRNPK (Shin *et al*. 2017; Fan *et al*. 2015) and its *C. elegans* homolog, HRPK-1 (Li *et al*. 2019) have been shown to impact miRNA-mediated gene regulation, albeit through potentially distinct mechanisms. hnRNPA2B1 was identified as a reader of m6A RNA modification that binds to m6A modified primary miRNAs, among other RNAs, and recruits the protein complex responsible for miRNA processing in the nucleus (Alarco’ *et al*. 2015). Sumoylated hnRNPA2B1 regulates sorting of miRNAs into exosomes, potentially influencing miRNA mediated cell-cell signaling (Villarroya-Beltri *et al*. 2013). hnRNPA1 has also been reported to regulate processing of primary miRNAs pri-miR-18a and pri-let-7 (Guil and Ca’ceres, 2007; Michlewski and Ca’ceres, 2008; Kooshapaur *et al*. 2018). While depletion of hnRNPA1 led to reduction in the levels of miR-18, let-7 levels were found to increase in Hela cells lacking hnRNPA1 (Guil and Ca’ceres, 2007; Michlewski and Ca’ceres, 2008; Kooshapaur *et al*. 2018). However, since changes in mature miRNA levels do not always correlate with changes in miRNA function *in vivo*, further characterization of hnRNPA1 and its homologs’ roles in gene regulatory events is needed.

We previously identified HRPA-1 in 2’-O methyl oligonucleotide-mediated let-7 and miR-58 pulldowns (Hebbar *et al*. 2022). HRPA-1 was previously reported to be a homolog of hnRNPA1 (Jeong *et al*. 2004), however, recent reports suggest HRPA-1 to be a closer homolog of hnRNPA2B1 (Ryan and Hart, 2021; Ryan *et al*. 2021). Like its human homologs, HRPA-1 has been shown to regulate telomere ends (Jeong *et al*. 2004) and alternative splicing (Barberan-Soler *et al*. 2011). Dysregulation of both human and *C. elegans* hnRNPA/B proteins have been implicated in neurodegeneration (Ryan and Hart, 2021; Ryan *et al*. 2021; Clarke *et al*. 2021). Another key feature that appears to be conserved among these proteins is their ability to form phase separations *in vitro* (Ritsch *et al*. 2022; Ryan *et al*. 2021). HRPA-1 has also been reported to form aggregates under stress in specific tissues when expressed from tissue specific promoters (Lechler and David, 2017; Ryan *et al*. 2021). The relatively high degree of sequence similarity between the hnRNPA/B proteins and HRPA-1 makes *C. elegans* a great system to evaluate hnRNP roles in developmental processes *in vivo*. In this study, we report our investigation into the role of HRPA-1 in miRNA-mediated gene regulation and the associated developmental processes. We found that loss of *hrpa-1* enhanced developmental defects of multiple reduction-of-function miRNA mutants, dysregulated mature miRNAs levels, and disrupted gene expression. Overall, we propose several models describing how HRPA-1 may contribute to miRNA-mediated gene regulation during *C. elegans* development.

## Materials and Methods

### Strain maintenance

All *C. elegans* strains were maintained on NGM and fed with *E. coli* strain OP50. Strains were maintained at 20°C unless otherwise noted.

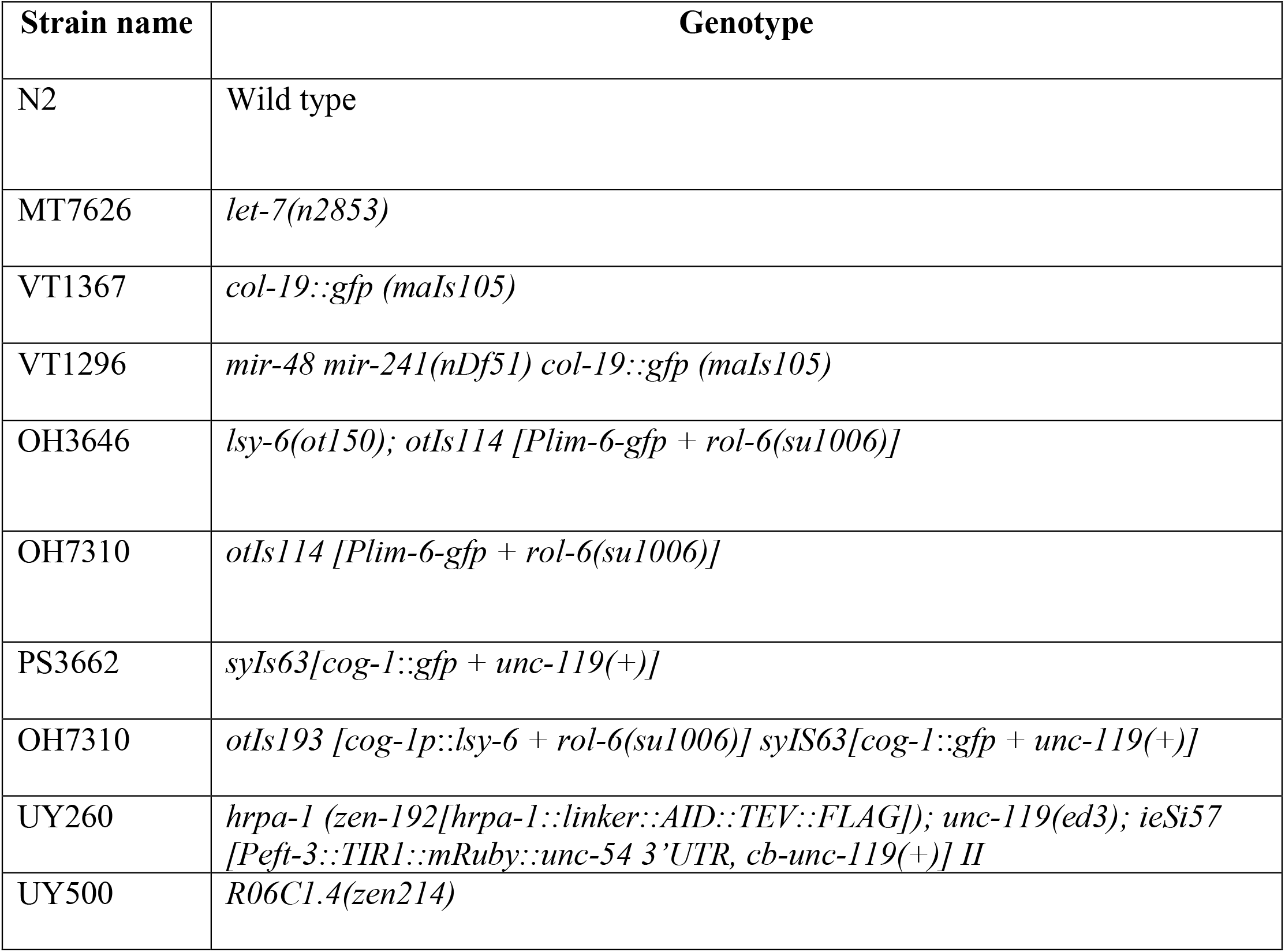
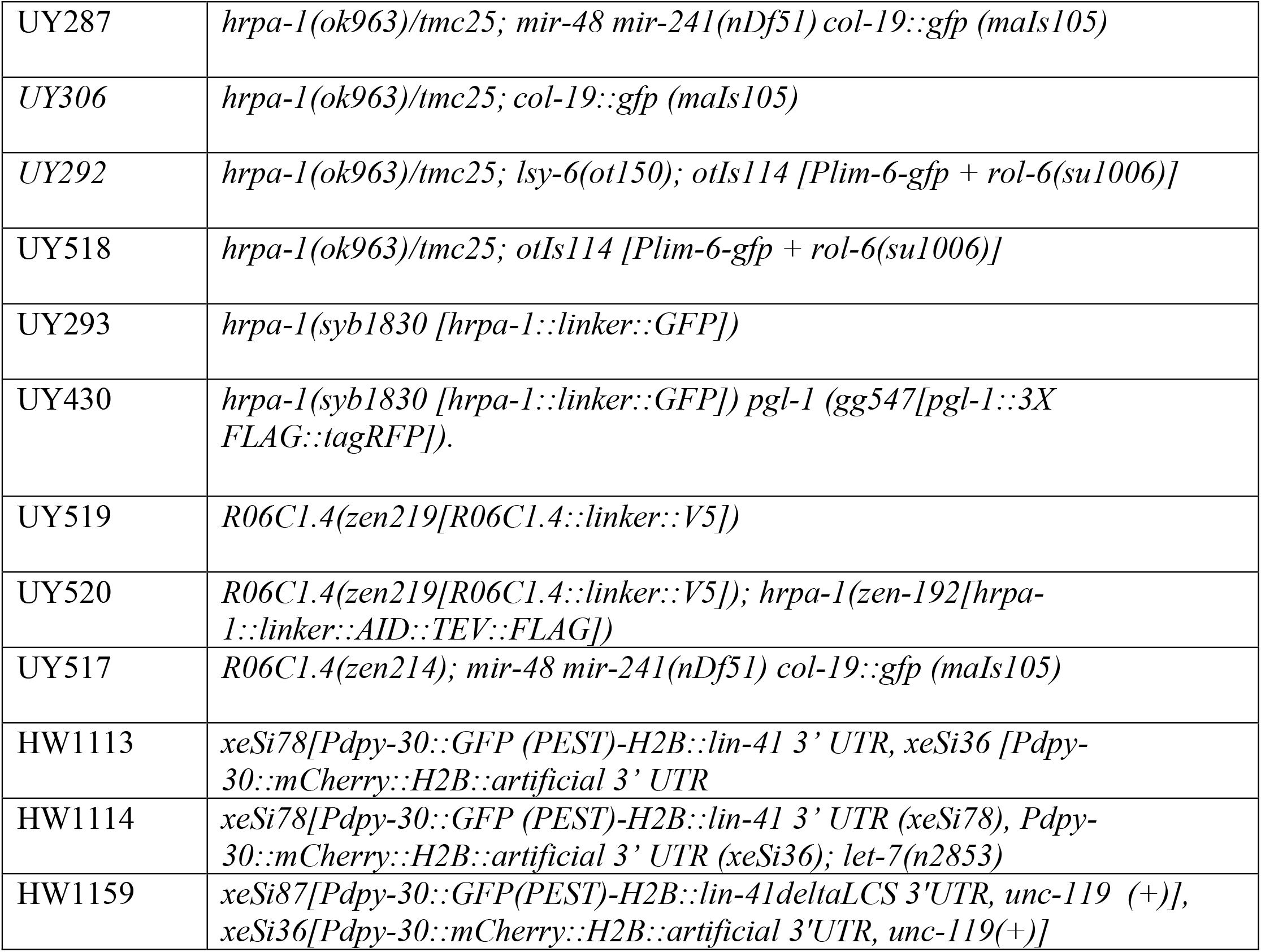

### RNAi and phenotypic assessments

For all phenotypic assays using *hrpa-1(ok963)* mutants, *hrpa-1(ok963)/hrpa-1(ok963)* progeny of *hrpa-1(ok963)/tmc25* parent in wild type or miRNA mutant backgrounds *(lsy-6(ot150)* or *mir-48 mir241(nDf51))* were scored. For ASEL cell fate assay, homozygous *hrpa-1(ok963); lsy-6 (ot150)* animals lacking the *plim-6::gfp* reporter expression in ASEL neurons were scored as “cell fate defective”. RNAi knockdown was performed as previously described (Kamath *et al*. 2003). Empty vector and *dcr-1* RNAi were used as negative and positive controls respectively. For the let-7 vulval bursting assay, N2 or *let-7(n2853)* embryos obtained through bleaching were placed on RNAi plates and grown at 15°C. Vulval bursting was scored in day 1 adults. For *cog-1* assay, *cog-1::gfp* or *pcog-1::lsy-6; cog-1::gfp* animals were transferred as L1s on RNAi plates and F1 progeny were scored at L4 stage for *cog-1::gfp* expression in uterine cells. Animals lacking reporter expression in one or both uterine cells were scored as abnormal.

For RNAi based assay, *mir-48 mir-241(nDf51) col-19::gfp (maIs105)* or *col-19::gfp (maIs105)* animals were placed on RNAi plates at L3 stage and the hypodermal and seam cell *col-19::gfp* expression were scored in F1 young adult animals. Seam cell numbers were scored by counting the number of seam cells between the pharynx and the anus.

### CRISPR-based genome editing

CRISPR/Cas9 based gene editing was performed by carrying out microinjections as previously described (Mello *et al*. 1991). Strains were generated by injecting RNP injection mix consisting of Alt-R Cas9 (IDT, cat# 1081058), *dpy-10* crRNA, gene specific crRNAs, and tracer RNA (IDT, cat# 1072532). Endogenously tagged *R06C1*.*4::V5* was generated by inserting coding sequences of V5 tag into the C-terminus of *R06C1*.*4* locus just before the stop codon through CRISPR/Cas9 mediated homologous recombination. *GFP* insertion at the endogenous *hrpa-1* locus to generate *hrpa-1 hrpa-1::linker::GFP* was done by SunyBiotech.

### HRPA-1 depletion, RNA extraction, Small RNA and RNA seq

HRPA-1 was depleted using RNAi and auxin treatment. Indole-3-acetic acid (IAA) was purchased from ThermoFisher (Product #I3750). IAA treatment was performed as previously described (Zhang *et al*. 2015). For HRPA-1 depletion, *hrpa-1::linker::AID*::TEV::FLAG* embryos obtained through bleaching were transferred to empty vector control or *hrpa-1* RNAi plates and maintained at 20ºC until the L4 stage. RNAi-treated L4 larvae were then transferred to either control or 1mM IAA plates for 3 hours before collecting the treated animals for downstream experiments. Western blot experiments after RNAi and IAA treatment were performed by picking 50 worms directly into the 2x Laemmli protein loading buffer and boiled at 95°C for 5 minutes.

RNA from L4 staged animals from control and *hrpa-1-*depleted conditions were extracted as previously described (Li *et al*. 2019), except worm pellets resuspended in Trizol were vortexed for 10 mins before proceeding to phenol-chloroform extraction step. Small RNA and RNAseq was performed on the same set of RNA. For small RNA library preparation, small RNA size selection was performed as previously described (Gu *et al*. 2011). NEXTflex Small RNA Library Prep kit v3 (Bioo Scientific) was used to prepare libraries according to the manufacturer’s instructions, followed by size selection of final PCR products as previously described (Gu *et al*. 2011). TruSeq stranded RNA-seq library prep was employed to prepare RNA libraries. All libraries were sequenced using the Illumina Nextseq500 platform at the Kansas State Genomics Core.

### Small RNA data analysis

Small RNAseq reads were checked for quality before and after filtering using FastQC v0.11.8 (https://www.bioinformatics.babraham.ac.uk/projects/fastqc). Cutadapt tool was used to: (1) clip the adapter sequence from 3’ end (-a ATCTCGTATGCCGTCTTCTGCTTG -e 0.1); (2) split reads into libraries using fastx_barcode splitter utility (http://hannonlab.cshl.edu/fastx_toolkit/index.html) based on their barcode index sequences and concatenate files that belong to the same barcode; (3) clip the remaining 3’ end and 5’ adapter sequences (-a TGGAATTCTCGGGTGCCAAGGAACTCCAGTCAC -e 0.1); (4) trim the first and last 4 random bases, and (5) select reads with a final length ranging between 17-29 nt for further analysis. Reads were mapped to *C. elegans* genome (WS279) using bowtie v1.2.2 (Langmead *et al*. 2009) allowing three mismatches in the alignment. Mature miRNA expression was quantified using the miRDeep2 pipeline (Friedländer *et al*. 2012). The DESeq2 package in R was used to perform differential expression analysis (Anders and Huber, 2010).

### RNAseq read processing and data analysis

FastQC v0.11.8 was used to assess quality of paired end RNA seq reads. Trimmomatic tool was used for quality filtering and adapter trimming. Filtered reads were mapped to *C. elegans* reference genome (WS279) using STAR aligner (Doblin *et al*. 2013). Stringtie transcript assembler was used to construct scaffold and contigs to assemble the transcripts. Normalized expression for each gene in the assembled transcriptome was quantified using featureCounts program (Subread package) (Liao *et al*. 2014). DESeq2 package in R was used to perform differential expression analysis (Anders and Huber, 2010).

### Protein extract preparation, miRNA pulldown and immunoprecipitation experiments

Protein extracts from mixed stage animals were prepared as previously described (Li and Zinovyeva, 2020). 2’O-methylated oligonucleotide-mediated miRNA pulldown was performed as previously described, using 1mg total protein for each pulldown (Jannot *et al*. 2011). HRPA-1::FLAG immunoprecipitation was performed using the anti-FLAG antibody-conjugated ChromoTek DYKDDDDK Fab-Trap™ Agarose beads (cat# ffa-10) according to the manufacturer’s protocol. Lysis and wash buffers for these immunoprecipitations were prepared as previously described (Li and Zinovyeva, 2020). Lysates with equal amounts of total protein from N2 and *hrpa-1::AID*::TEV::FLAG* strains were used for FLAG IP. Immunoprecipitation was performed for 1h at 4°C followed by 4 washes with the Wash buffer. 10% of the input was used for western blot. HRPA-1::AID*::TEV::FLAG was detected using Monoclonal Anti-FLAG M2-Peroxidase (HRP) antibody (A8592-2MG) from Sigma-Aldrich at a 1:1000 dilution. R06C1.4::V5 was detected using the Anti-V5 antibody from Invitrogen (MA5-32053) at a 1:1000 dilution. ALG-1 was detected using custom generated anti-ALG-1 monoclonal antibody (Zinovyeva *et al*. 2014). As a loading control, mouse Anti-TUBULIN antibody from Sigma-Aldrich (T6074) was used at a 1:5000 dilution to detect tubulin.

FLAG-based RNA immunoprecipitation experiments were performed using lysates from N2 and *hrpa-1::AID*::TEV::FLAG* strain. Lysates with equal amount of total protein was used across both samples for immunoprecipitation, RNA extraction, and western blots. Following immunoprecipitation, 25% of the suspension was used for western blot and the rest was used for total RNA extraction using the phenol/chloroform method.

### Western blot detection of HRPA-1 from mutants

Western blot analyses for detection of HRPA-1 from *hrpa-1(ok963)* and *hrpa-1(ok597)* mutants were performed by directly picking ∼50 worms into 2x Laemmli protein loading buffer, with custom Rabbit polyclonal anti-HRPA-1 antibody (Jeong *et al*. 2004) used for detection.

### RT-qPCR

RNA quantification following FLAG IP was performed by RT-qPCR using the one-step QuantiNova SYBR-Green RT-PCR kit (cat# 208152) according to the manufacturer’s protocol. ABI 7500 Real-Time PCR System was used to perform all the RT-qPCR experiments. Ct values were adjusted to account for input and IP RNA dilutions. To calculate % input, we first normalized the Ct value for IP RNAs with their respective input RNAs to obtain Δ Ct. Fold enrichment was determined by Δ Δ Ct method by further normalizing *hrpa-1::AID*::TEV::FLAG* RIP fraction to our negative control (IP fraction from N2).

### Microscopy and statistical analyses

Phenotypes were scored using a fluorescence-equipped Leica DM6 upright microscope and images were captured using Leica DM6B camera. Images were processed using the Leica Application Suite X (3.4.1.17822) software. HRPA-1::GFP images were assembled using Adobe Illustrator. Student t-test was used to determine statistical significance for RNAi based assays (hypodermal *col-19::gfp*, seam cell number analysis, vulval bursting assay and *cog-1* reporter assay). Chi-square test was performed to determine statistical significance for all other assays. GraphPad Prism (9.2.0 (332)) software was used to perform all of the statistical analyses.

## Results

We previously characterized miRNA-interacting protein complexes to identify factors that coordinate with miRNAs to regulate gene expression (Hebbar *et al*. 2022). HRPA-1, an RNA binding protein, was captured in 2-O’methylated oligonucleotide mediated pulldowns of both let-7 and miR-58 miRNAs (Fig 1A, Hebbar *et al*. 2022). To begin to understand the role of HRPA-1 in miRNA mediated gene regulation, we first sought to confirm the precipitation of HRPA-1 in 2’-O-methyl oligonucleotide-mediated miRNA pulldowns. let-7- and miR-58-specific oligos, but not the scrambled control oligo, precipitated FLAG-tagged HRPA-1 from an endogenously tagged *hrpa-1::AID*::TEV::FLAG* strain (Fig 1B). In addition, immunoprecipitation of HRPA-1::AID*::TEV::FLAG co-precipitated the miRISC component, ALG-1 (Fig 1C), consistent with previous detection of HRPA-1 in ALG-1 IP through mass-spec analysis, with average fold enrichment of 2.94 (Zinovyeva *et al*. 2015). The small amount of ALG-1 precipitating in HRPA-1::AID*::TEV::FLAG IP, as well as the mild enrichment observed in ALG-1 mass spec (Zinovyeva *et al*. 2015) suggest that HRPA-1 interaction with ALG-1 may be transient and may be RNA-dependent.

**Figure 1.**
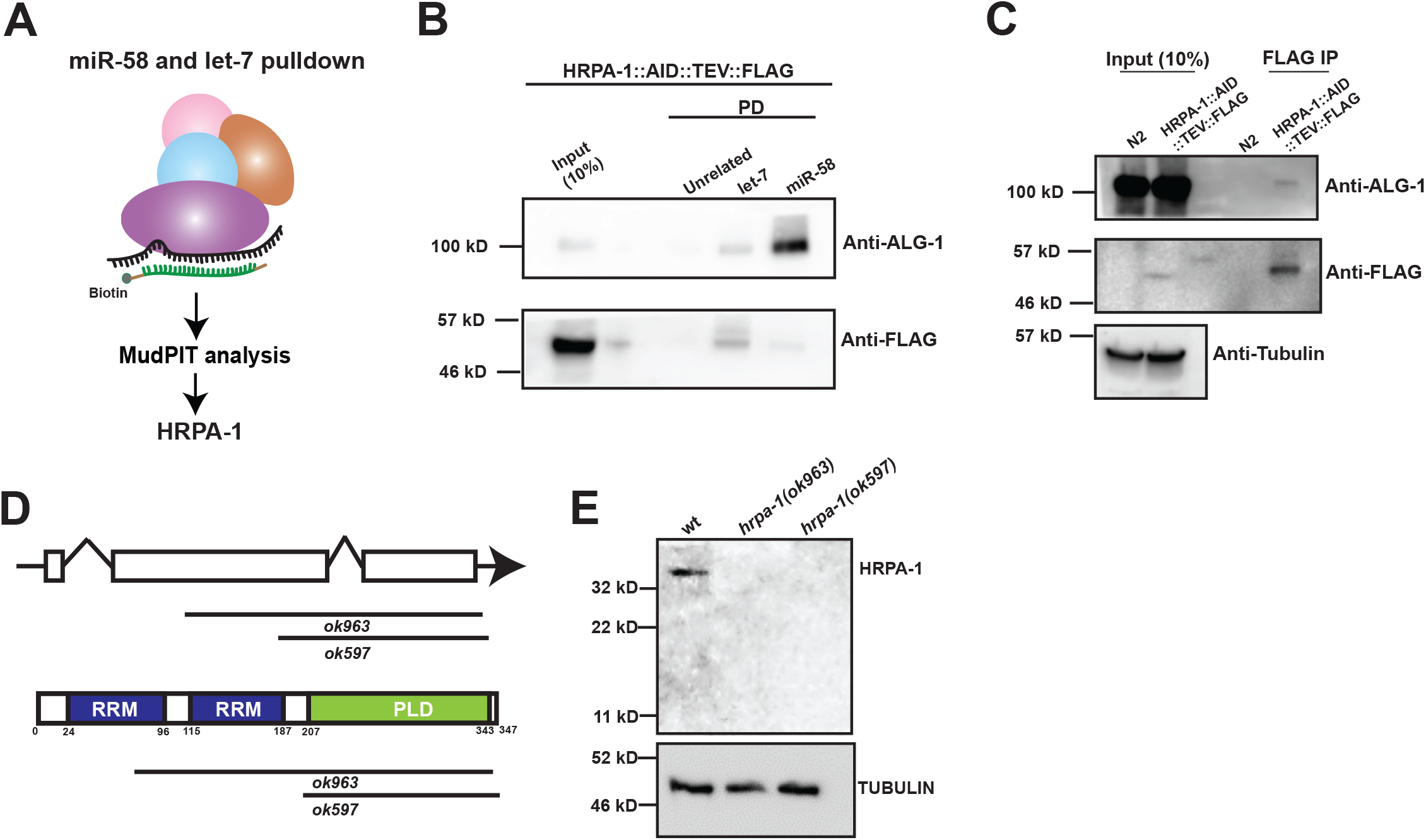
HRPA-1 was captured in 2’ O-methyl oligonucleotide-mediated let-7 and miR-58 pulldowns. (A) HRPA-1 was previously captured in let-7 and miR-58 pulldown experiments (Hebbar *et al*. 2022). (B) Detection of HRPA-1 and ALG-1 in Western blot following 2’-O-methylated oligonucleotide-mediated miRNA pulldowns (PD) from *hrpa-1::AID::TEV::FLAG* animals. (C) HRPA-1 immunoprecipitation co-precipitates ALG-1. (D) A schematic of *hrpa-1* gene structure, protein domains, and available *hrpa-1* deletion mutations. RRM-RNA Recognition Motif, PLD-Prion like Domain. E) *hrpa-1* mutants do not produce detectable protein by Western blot analysis.

To assess genetic interactions between *hrpa-1* and miRNAs of interest, we confirmed that the existing *hrpa-1* deletion mutants, *hrpa-1(ok963)* (Jeong *et al*. 2004) and *hrpa-1(ok597)* (Ryan *et al*. 2021) are strong loss of function/null based on the molecular nature of the alleles and the lack of detectable HRPA-1 protein (Fig 1D, E). For subsequent functional analyses, we used the larger of the two deletions, *hrpa-1(ok963)*.

### *hrpa-1* genetically interacted with let-7 family miRNAs

HRPA-1 co-precipitated in a let-7 complementary oligonucleotide pulldown, which captures multiple members of the *let-7* family of miRNAs (Zinovyeva *et al*. 2014). We wanted to determine whether *hrpa-1* is functionally important for the *let-7* family miRNAs. *let-7* family miRNAs, let-7, miR-48, miR-241, and miR-84 are integral components of the heterochronic pathway that controls the temporal progression of *C. elegans* post-embryonic developmental programs (Reinhart *et al*. 2000; Abbott *et al*. 2005). Highly conserved *let-7* plays crucial roles during *C. elegans* L4 larval to adult transition (Reinhart *et al*. 2000). In addition to governing hypodermal development, *let-7* also regulates vulval development, primarily by repressing *lin-41* in vulval precursor cells (Ecsedi *et al*. 2015). Loss of *let-7* leads to derepression of *lin-41*, which results in a burst vulva phenotype (Reinhart *et al*. 2000). A temperature sensitive *let-7(n2853)* mutant has a partially penetrant vulval bursting phenotype at permissive temperature (15ºC), making it amenable to genetic perturbations (Fig 2A, Vella *et al*. 2004). Knockdown of *hrpa-1* in *let-7(n2853)* mutant at 15°C enhanced the vulval bursting phenotype, suggesting that *hrpa-1* may directly or indirectly facilitate let-7 mediated target repression during vulval development (Fig 2B, Table S2).

**Figure 2.**
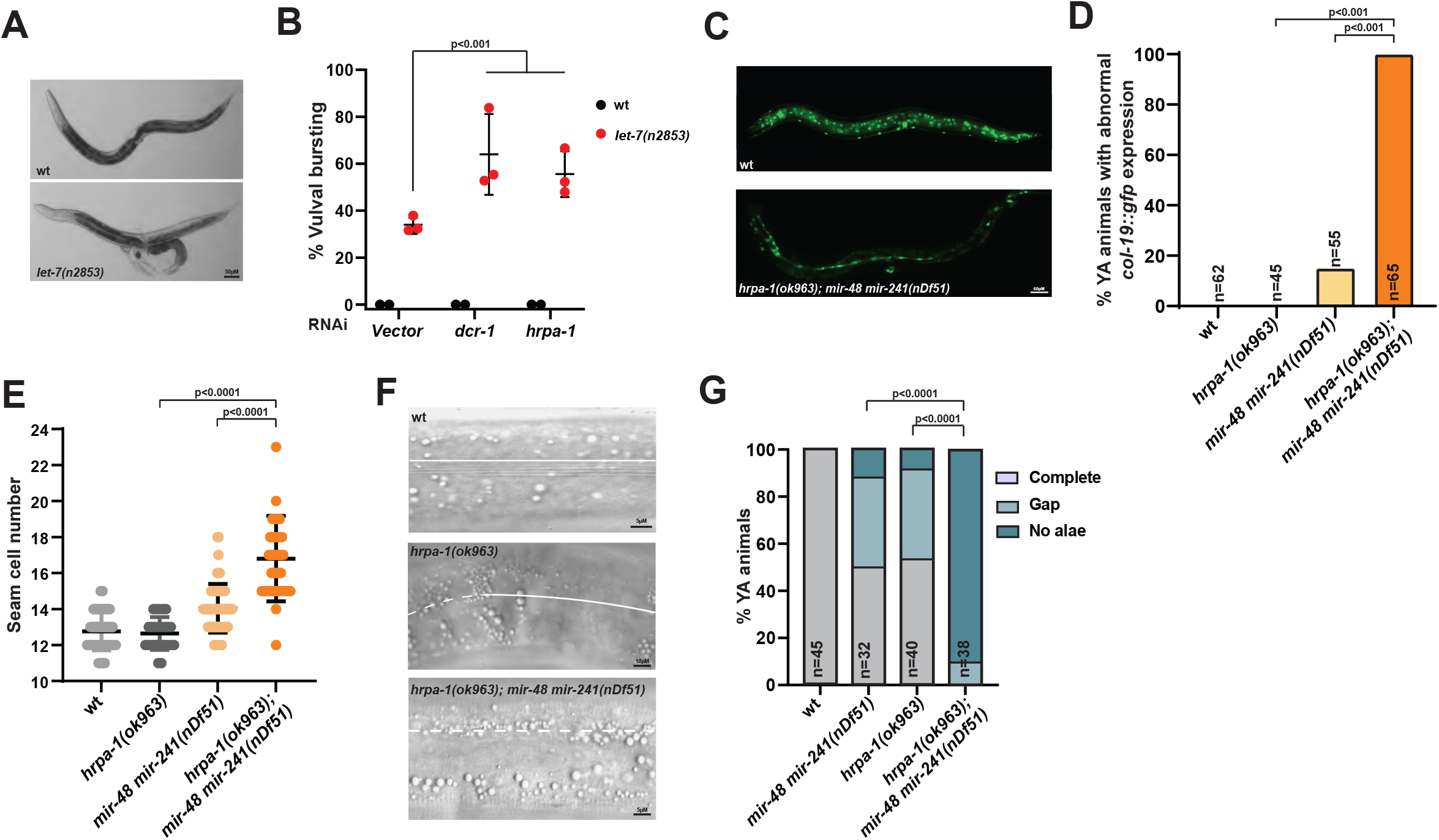
Loss of *hrpa-1* enhances developmental defects associated with *let-7* family miRNA mutants. (A) *let-7(n2853)* animals show a partially penetrant vulval bursting phenotype at 15ºC. (B) RNAi knockdown of *hrpa-1* enhances the vulval bursting phenotype at 15ºC. (C) *mir-48 mir-241(nDf51)* young adults show delayed hypodermal expression of *col-19::gfp* reporter. *hrpa-1(ok963)* enhances the (D) abnormal hypodermal *col-19::gfp* expression, (E) seam cell lineage defect, and (F-G) alae formation defect of *mir-48 mir-241(nDf51)* animals. Solid lines indicate alae, dashed lines indicate regions lacking discernable alae. T-test was used to determine the statistical significance for let-7 vulval bursting and seam cell number analyses. Chi-square was used to analyze hypodermal *col-19::gfp* and alae defect assays.

To evaluate the possibility of HRPA-1 regulating miRNA targets in a miRNA-independent manner, we determined the effect of *hrpa-1* knockdown on a well-established let-7 target, *lin-41*, whose dysregulation leads to vulval bursting (Ecsedi *et al*. 2015). A fluorescent reporter-based quantification of *lin-41_3’UTR* levels (Ecsedi *et al*. 2015) in wild type and *let-7(n2853)* at non-permissive temperature (20ºC) did not show any changes in reporter level upon *hrpa-1* knockdown (Fig S1, Table S2). Further, *lin-41deltaLCS 3’UTR* reporter, which harbors let-7 complementary site deletion, showed no changes in reporter levels upon *hrpa-1* knockdown (Fig S1, Table S2). These observations suggest that *hrpa-1* might not regulate *lin-41* on its own and the enhancement of vulval bursting in *let-7(n2853)* assay may be due to derepression of let-7-mediated *lin-41* repression.

miR-48, miR-241 and miR-84 redundantly target *hbl-1* for repression during L2-L3 transition, ensuring the timely division and differentiation patterns of *C. elegans* seam cells (Abbott *et al*. 2005). Complete loss of these miRNAs leads to reiteration of L2 specific cell divisions and delayed expression of the adult-specific marker *col-19::gfp* (Abbott *et al*. 2005). Partial deletion mutant *mir-48 mir-241(nDf51)* shows incompletely penetrant delayed development of the hypodermal tissue, as assayed by *col-19::gfp* expression in the seam cells and the hypodermis (Fig 2C, Abbott *et al*. 2005). Deletion of *hrpa-1* in this background enhanced the heterochronic defects of *mir-48 mir-241(nDf51)* mutant (Fig 2D, E, Table S3), observed through delayed *col-19::gfp* expression in hypodermal cells (Fig 2D) and increased seam cell numbers in young adult animals (Fig 2E). Loss of *hrpa-1* alone did not delay hypodermal development or affect seam cell number (Fig 2D, E, Table S3). However, *hrpa-1(ok963)* mutants had alae formation defects (Fig 2F, G, Table S3). Deleting *hrpa-1* in *mir-48 mir-241(nDf51)* background further enhanced these alae formation defects (Fig 2F, G, Table S3), suggesting that *hrpa-1* is involved in developmental processes controlled by the *let-7* family miRNAs.

### Loss of *hrpa-1* enhanced developmental defect associated with *lsy-6(ot150)*

To determine whether *hrpa-1* broadly coordinates with miRNAs to regulate *C. elegans* development, we examined the effects of *hrpa-1(ok963)* on *lsy-6* miRNA activity. *lsy-6* functions to specify asymmetric cell fates of chemosensory neurons. *lsy-6* specifies ASEL cell fate _by_ repressing *cog-1* which encodes an ASER promoting transcription factor (Johnston and Hobert, 2003). Loss of lsy-6 activity results in derepression of *cog-1* leading to defective specification of ASEL cell fate, which can be tracked using a *plim-6::gfp* reporter (Johnston and Hobert, 2003). A hypomorphic mutant, *lsy-6(ot150)*, shows a partially penetrant cell fate defective phenotype (Fig 3A, B). *hrpa-1(ok963)* enhanced the *lsy-6(ot150)* cell defective phenotype but did not produce an ASE cell fate defect on its own (Fig 3B, Table S3). To determine whether *hrpa-1* is important for lsy-6 mediated *cog-1* repression, we utilized the *cog-1* reporter strains (Palmer *et al*. 2002). *cog-1* is normally expressed in the vulval and uterine cells (Palmer *et al*. 2002). Uterine *cog-1::gfp* expression is repressed in the reporter strain that expresses *lsy-6* under *cog-1* promoter (Fig 3C, Palmer *et al*. 2002). Knockdown of *hrpa-1* alone did not affect *cog-1::gfp* expression (Fig 3D, Table S2). However, RNAi of *hrpa-1* in the *pcog-1::lsy-6; cog-1::gfp* reporter strain derepressed *cog-1::gfp* in the uterine cells (Fig 3D, Table S2), suggesting that *hrpa-1* may be important for lsy-6-mediated *cog-1* repression in that tissue.

**Figure 3.**
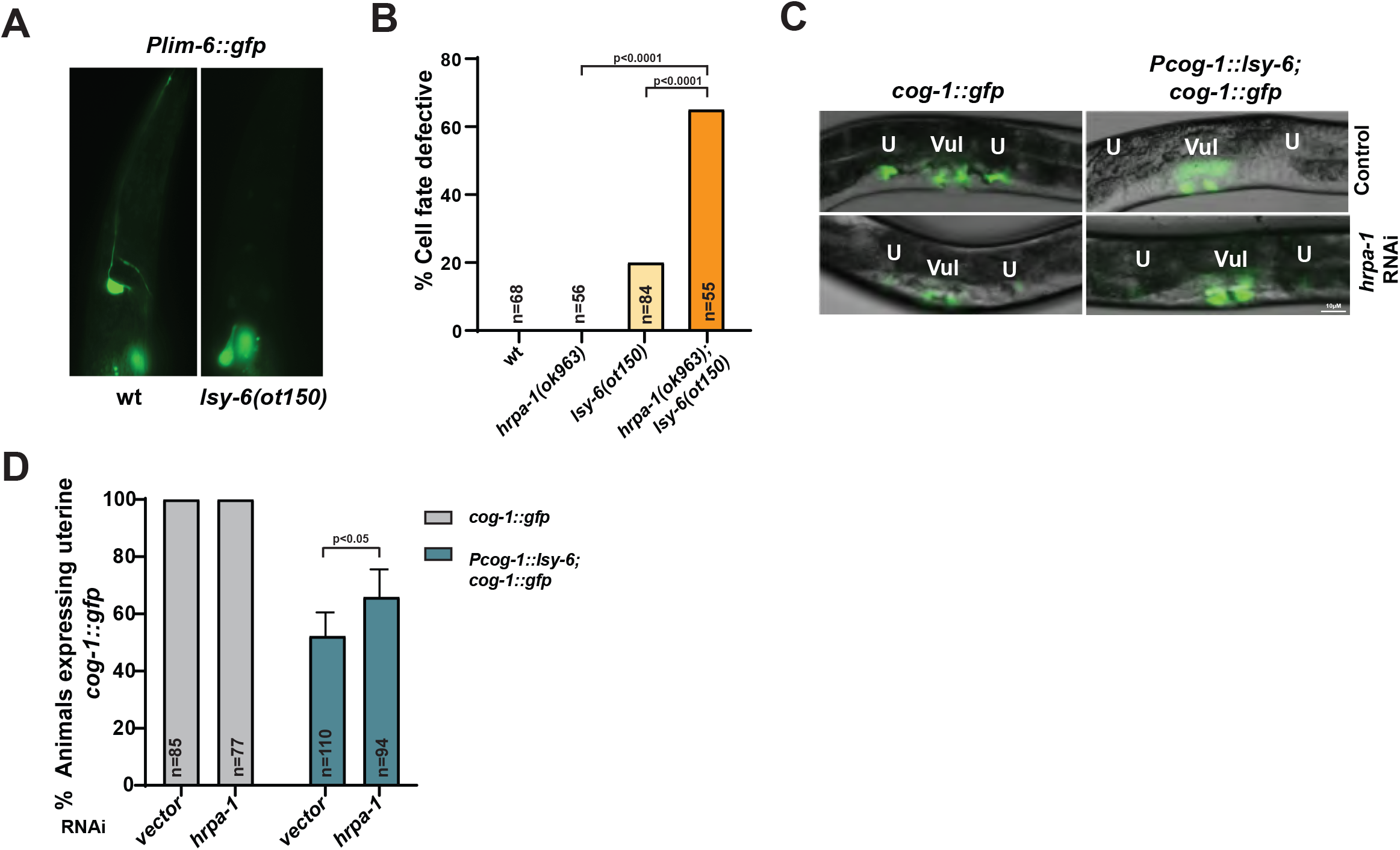
*hrpa-1* is functionally important for lsy-6 miRNA activity. (A) *lsy-6(ot150)* mutants show partially penetrant defective ASEL neuronal cell fate phenotype. (B) *hrpa-1(ok963)* enhanced the ASEL cell defective phenotype of *lsy-6(ot150)* animals. (C-D) *cog-1* expression in the uterine cells is repressed when *lsy-6* is expressed in the same tissue. Depletion of *hrpa-1* leads to derepression of lsy-6 mediated *cog-1* repression in the uterine cells (C), quantified in (D). Chi-square and T-test were used to determine the statistical significance of ASEL cell fate assay and *cog-1* reporter assay respectively.

### HRPA-1 is ubiquitously expressed

To determine the spatial and temporal expression pattern of HRPA-1, we tagged the 3’ end of the endogenous *hrpa-1* gene with GFP. HRPA-1::GFP was strongly nuclear and showed ubiquitous expression across somatic and germline tissues throughout *C. elegans* development (Fig 4A-C). Weak cytoplasmic GFP signal was detected in the intestine (Fig 4C). Our finding is consistent with a previous report describing nuclear expression of HRPA-1 assessed by a multicopy extrachromosomal *hrpa-1* transgene (Joeng *et al*. 2004). RFP/mCherry tagged *hrpa-1* transgenes expressed under tissue specific promoters have been reported to form cytoplasmic puncta in pharynx and aggregates in neurons under stress (Lechler and David, 2017; Ryan *et al*. 2021). To determine whether endogenous promoter driven HRPA-1 also forms aggregates under stress, we subjected *hrpa-1::gfp* animals to heat stress at 35ºC for 30 min. Heat stress induced HRPA-1::GFP localization to the perinuclear region forming puncta which overlapped with PGL-1::tagRFP signals (Fig 4D, D’), a key component of P-granules crucial for post transcriptional gene regulation.

**Figure 4.**
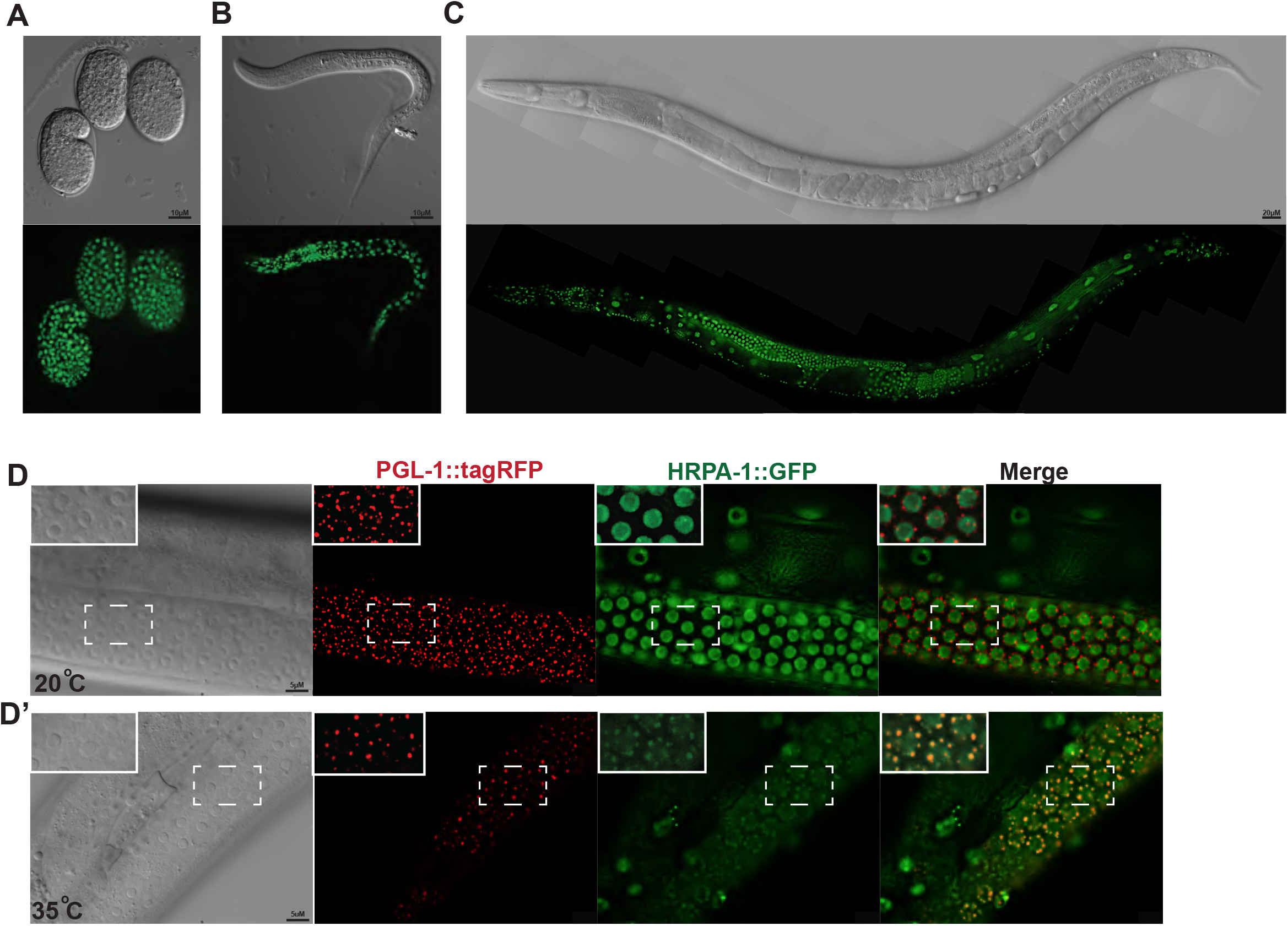
HRPA-1::GFP is expressed in the nucleus and localizes to the germline perinuclear region upon heat stress. Endogenously tagged HRPA-1::GFP expression in (A) embryos, (B) L2 stage larvae, and (C) adults. HRPA-1::GFP is strongly expressed in the nucleus in somatic tissues and in the germline. (D, D’) HRPA-1::GFP expression pattern in the germline at 20ºC and (D’) at 35ºC. HRPA-1::GFP localizes to the perinuclear region upon heat shock, colocalizing with PGL-1::tagRFP (D’).

### Depletion of *hrpa-1* altered miRNAs abundance profiles and dysregulated gene expression

A functional requirement of *hrpa-1* in miRNA-mediated gene regulation could stem from HRPA-1’s role in miRNA biogenesis and/or miRNA activity. To determine HRPA-1 involvement in miRNAs biogenesis, we assessed levels of mature miRNAs following HRPA-1 depletion. *hrpa-1(-/-)* animals show severe developmental defects including embryonic lethality, reduced brood size, and larval arrest (Barberan-Soler *et al*. 2011; Ryan and Hart, 2021, Hills-Muckey *et al*. 2022). Thus, to deplete HRPA-1, we subjected *hrpa-1::AID*::TEV::FLAG* animals to both RNAi treatment and IAA induced protein depletion (Fig 5A), since neither knockdown alone resulted in robust depletion (Fig 5B,C). Small RNAseq from HRPA-1-depleted L4 animals revealed that expression levels of most miRNAs were unaffected by loss of HRPA-1 (Fig 5D). Expression levels of 19 miRNAs were disrupted by loss of HRPA-1, with the levels of 11 miRNAs upregulated (ex: miR-320, miR-243) and 8 downregulated (ex: miR-1817), suggesting that HRPA-1 function may be required for biogenesis or stability of these miRNAs (Fig 5D, Table S4). Levels of let-7 and miR-58 miRNAs were unaffected (Fig 5D, Table S4).

**Figure 5.**
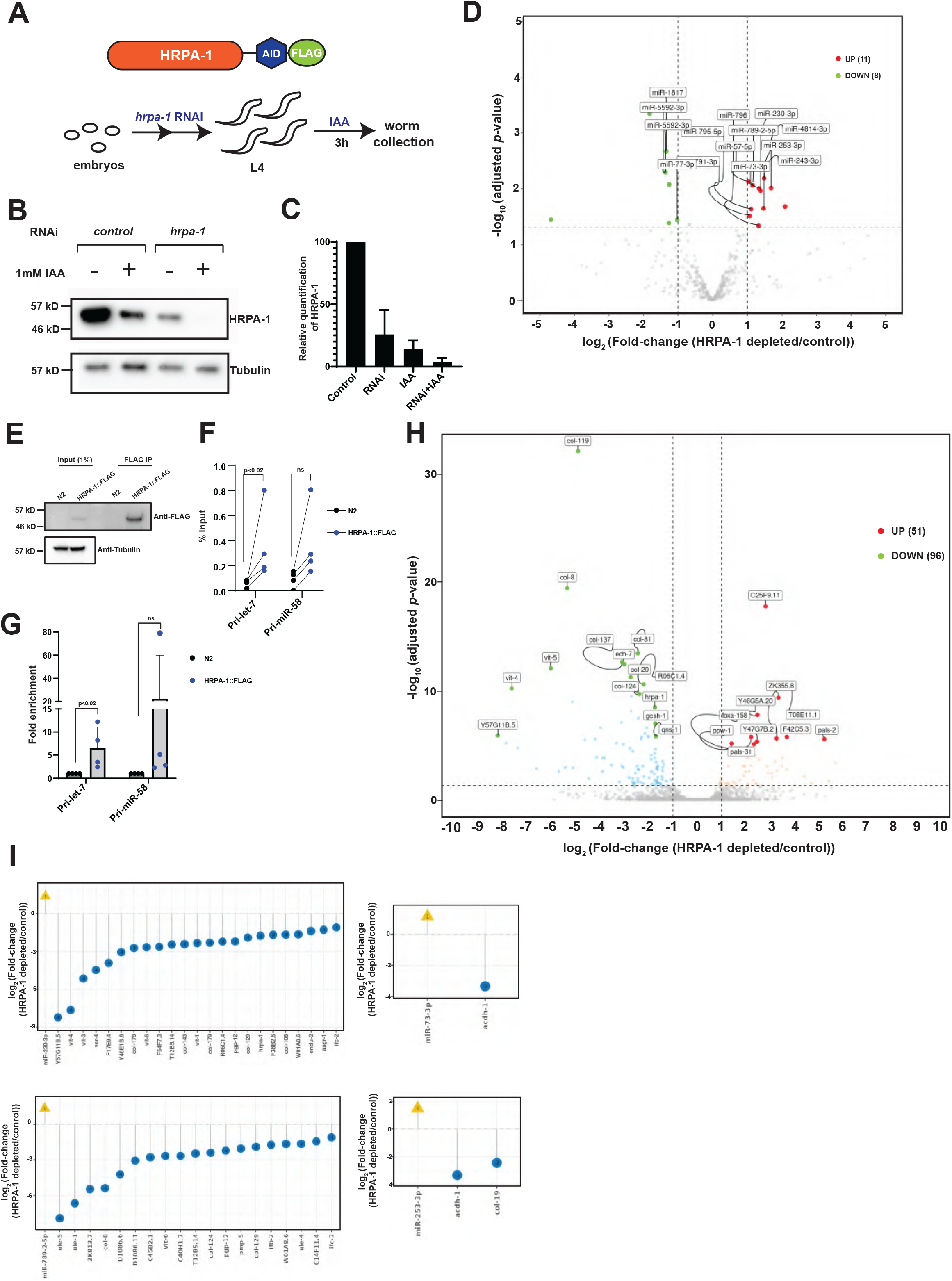
HRPA-1 depletion disrupts mature miRNA and gene expression levels. (A-C) HRPA-1 was depleted using combined RNAi and IAA treatment from *hrpa-1::AID*::TEV::FLAG* animals. (A) A schematic of RNAi and IAA treatment performed to deplete HRPA-1. (B) Representative western blot image and (C) quantification of protein depletion achieved in the respective treatment conditions. (D) HRPA-1 depletion alters miRNA abundances as assayed by small RNAseq analysis. (E-G) Anti-FLAG immunoprecipitation from *hrpa-1::AID*::TEV::FLAG* or N2 (control) animals (E), followed by RT-qPCR quantification of bound pri-let-7 and pri-miR-58 RNA, shown as percent input (F) or fold enrichment over control (G). Paired T-test was used to determine statistical significance. (H) Depletion of HRPA-1 led to a disruption of gene expression as assayed by RNAseq analysis. (I) Some of the differentially expressed genes are predicted targets of miRNAs whose levels were disrupted by loss of HRPA-1. Lollipop plots showing log_2_(fold) changes in expression levels of miRNAs and their predicted targets, whose expression was altered in HRPA-1-depleted samples.

One of the human homologs of HRPA-1, hnRNPA1, associates with primary miRNAs *in vitro*. Since HRPA-1 was identified in miRNA complementary pulldown experiments that could, in theory, capture primary miRNAs, we sought to determine whether HRPA-1 IP may precipitate *let-7* and *mir-58* primary miRNAs. We performed an anti-FLAG based immunoprecipitation using *hrpa-1::AID*::TEV::FLAG* and N2 (negative control) lysates (Fig 5E). Real-time qPCR quantification of precipitated RNA showed a relative enrichment for pri-let-7 in HRPA-1 IP over our negative control (Fig 5F, G), suggesting that HRPA-1 may associate with pri-let-7. Pri-miR-58 was enriched, albeit not to statistically significant levels due to high variation (Fig 5F,G).

To gain insight into the effects of *hrpa-1* depletion on global gene expression, we performed RNAseq on the RNA extracted from genotype *hrpa-1::AID*::TEV::FLAG* L4 animals treated with *hrpa-1* RNAi and auxin (Fig 5A). 147 genes were differentially expressed in HRPA-1-depleted animals, with 96 genes downregulated and 51 genes upregulated (Fig 5H, Table S5). Since intronic miRNA transcription depends on their host gene and the splicing machinery (Kim and Kim, 2007), we wondered whether the changes in levels of intronic miRNAs were a result of changes in host gene transcription/processing. Expression levels of genes hosting intronic miRNAs were not affected by depletion of HRPA-1 (Fig S2, Table S4, S5), suggesting that the modest changes observed in intronic miRNA levels were not due to effects of *hrpa-1* on host gene transcription.

Since disruption of miRNA-mediated target repression could be the underlying cause of differential expression of some of the genes following depletion of HRPA-1, we asked whether genes disrupted upon loss of HRPA-1 were predicted targets of miRNAs affected by depletion of HRPA-1 (Fig 5I). We found that some of the predicted targets of upregulated miRNAs (such as miR-230, miR-243, miR-253, and miR-789-2) were downregulated by loss of HRPA-1 (Fig 5I). These targets included collagens (*col-19, col-8*, and *col-124*, among others) and other genes such as *R06C1*.*4, clec-4, acdh-1*, and *pmp-5* (Fig 5I).

### *R06C1*.*4* acts downstream of *hrpa-1* and depletion of *R06C1.4* partially recapitulates *hrpa-1* loss-of-function phenotypes in *mir-48 mir-241(nDf51)* background

Genes dysregulated by depletion of HRPA-1 could be the underlying cause of the enhanced developmental defects observed in miRNA mutant backgrounds upon loss of *hrpa-1*. To test this, we performed RNAi knockdown of top genes downregulated upon *hrpa-1* depletion (excluding collagen and vitellogenin genes). While *R06C1*.*4* and *ego-1* RNAi on their own did not result in developmental timing defects (Fig 6A, B), depletion of *R06C1*.*4* and *ego-1* in *mir-48 mir-241(nDf51)* background partially recapitulated the enhanced the developmental timing phenotype observed in *mir-48 mir-241; hrpa-1* animals (Fig 6A, B, Table S2). Knockdown of either of these genes did not alter the vulval bursting phenotype in *let-7(n2853)* assay at 15°C (Fig S3). To confirm the *R06C1*.*4* RNAi knockdown phenotype, we used CRISPR/Cas9 to delete the *R06C1*.*4* genomic locus (Fig 6C). *R06C1*.*4(zen214)* enhanced delayed hypodermal *col-19::gfp* expression and increased the seam cell number observed in *mir-48 mir-241* young adults (Fig 6D, E, Table S3), confirming the *R06C1*.*4* RNAi effects in *mir-48 mir-241(nDf51)* background (Fig 6A, B, Table S3).

**Figure 6.**
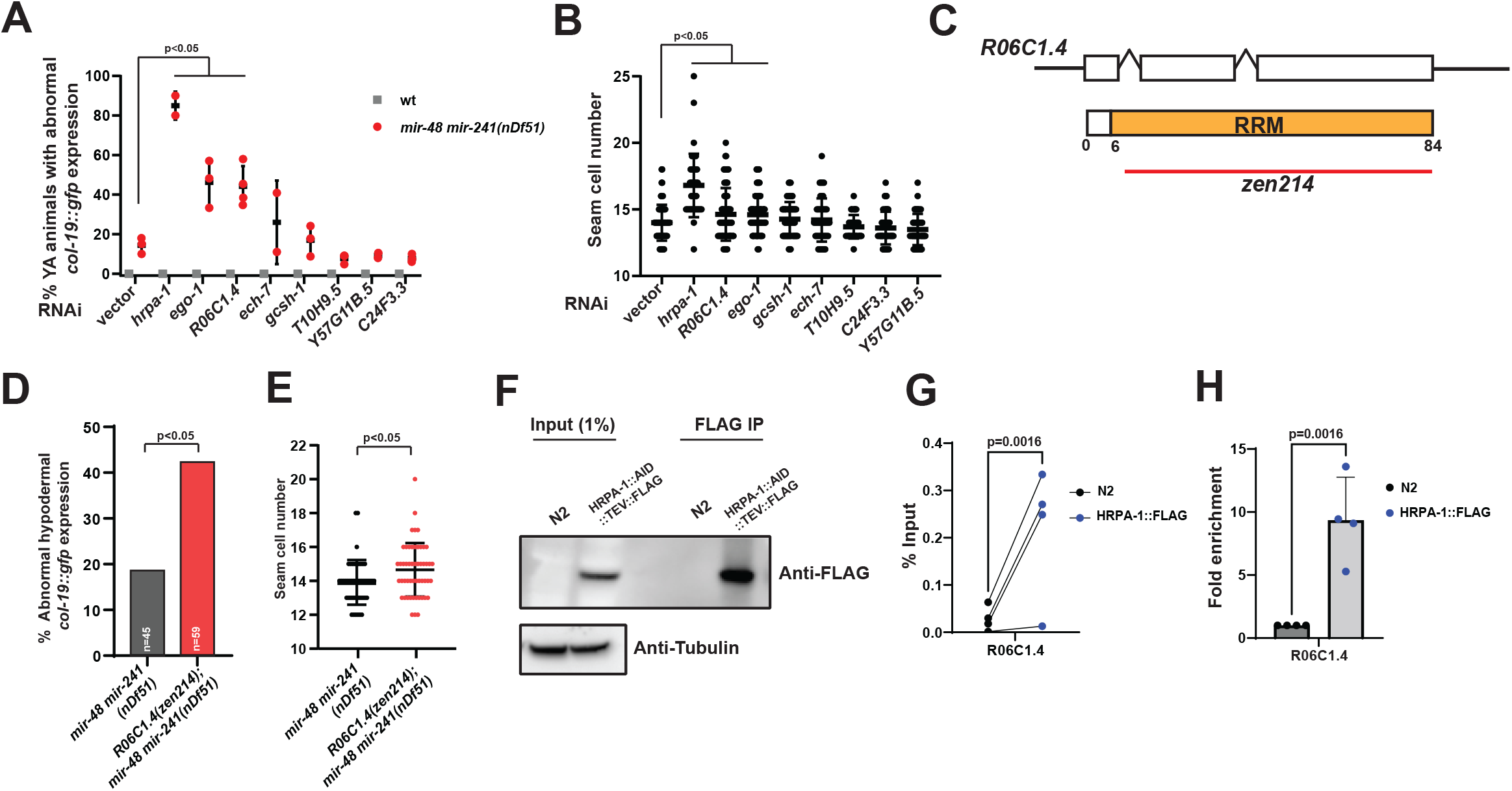
*R06C1*.*4* depletion partially recapitulates *hrpa-1* loss of function phenotype in *mir-48 mir-241(nDf51)* background. (A-B) RNAi knockdown of top non-collagen genes downregulated by loss of HRPA-1 in *mir-48 mir-241(nDf51)* background partially recapitulates (A) abnormal *col-19:gfp* expression and (B) seam cell number defect produced by *hrpa-1* depletion in *mir-48 mir-241* mutant animals. (C) *R06C1*.*4* gene and protein structure schematics. RRM–RNA Recognition Motif. Deletion allele, *R06C1*.*4(zen214)*, was generated using CRISPR/Cas9 genome editing deletes most of the *R06C1*.*4* gene. (D, E) *R06C1*.*4(zen214)* enhances the abnormal hypodermal *col-19::gfp* expression and seam cell lineage defect in *mir-48 mir-241(nDf51)* background. (F-H) Anti-FLAG immunoprecipitation from *hrpa-1::AID*::TEV::FLAG* or N2 (control) animals (E), followed by RT-qPCR quantification of bound R06C1.4 RNA, shown as percent input (F) or fold enrichment over control (G).

R06C1.4 appears to play a role in facilitating HRPA-1 function on miRNA-mediated developmental process. Since *R06C1*.*4* was downregulated upon *hrpa-1* depletion (Fig 5H), we wondered whether HRPA-1 could directly interact with *R06C1*.*4* mRNA. HRPA-1 IP significantly enriched for *R06C1*.*4* mRNA compared to N2 control, supporting a potential model for posttranscriptional regulation of *R06C1*.*4* RNA by HRPA-1 (Fig 6 G,H). The yeast homolog of R06C1.4, Rna15, is a component of cleavage and polyadenylation factor I (CFI) that is involved in 3’ end processing of precursor mRNAs (Fig S4A). Yeast hnRNP homolog, HRP1, is also a component of this complex (Gordon *et al*. 2011). To explore all avenues of HRPA-1 effects on *R06C1*.*4*, we wondered whether HRPA-1 could physically associate with R06C1.4 and perhaps be involved in 3’ end processing of primary miRNAs, in addition to feeding back onto *R06C1*.*4* expression. However, an anti-FLAG immunoprecipitation from *hrpa-1::AID*::TEV::FLAG* worms expressing R06C1.4::V5 did not co-precipitate R06C1.4 (Fig S4B). Thus, our data suggest that HRPA-1 does not form a physical complex with R06C1.4.

## Discussion

Heterogenous nuclear ribonucleoproteins have been implicated in several gene regulatory processes (reviewed in Geuens *et al*. 2016). Despite being extensively studied in different species, their complete functional profile is not entirely understood. The *C. elegans* hnRNPA/B family homolog, HRPA-1, has been previously reported to regulate telomere length, alternative splicing, and longevity (Jeong *et al*. 2004; Barberan-Soler *et al*. 2011). In this study, we report a role for *hrpa-1* in miRNA-regulated developmental processes.

We previously identified HRPA-1 in miR-58 and let-7 pulldowns (Hebbar *et al*. 2022) and we confirm the co-precipitation in this study (Fig 1B). While these data are suggestive of a direct interaction with mature let-7 and miR-58, we cannot rule out a potential interaction with primary or precursor miRNAs. Findings from HRPA-1 IP in this study (Fig 1C) and previously reported ALG-1 IP (Zinovyeva *et al*. 2015) indicate that HRPA-1 may transiently interact with ALG-1. The observed weak interaction between ALG-1 and HRPA-1 may be RNA-dependent, possibly stemming from HRPA-1 and miRISC binding to proximal binding sites on the targets. However, HRPA-1 was not previously identified in AIN-1 or AIN-2 immunoprecipitations (Zhang *et al*. 2007; Dallarie *et al*. 2018), suggesting HRPA-1 might not interact with mature miRISC, or may do so very transiently. Taken together, these observations indicate that HRPA-1 may interact with miRNA biogenesis intermediates prior to miRISC formation or may interact with miRISC, but transiently.

### Can subcellular HRPA-1 localization inform us of its function

The ubiquitous expression of HRPA-1::GFP suggests a role for this protein in multiple developmental processes (Fig 4A). A strong nuclear expression is consistent with the subcellular localization of its orthologs in humans, flies (Dreyfuss *et al*. 1993; Borah *et al*. 2008). The human homologs, hnRNPA1 and hnRNPA2/B1, have been reported to shuttle between nucleus and cytoplasm (Izaurralde *et al*. 1997). It’s possible that the cytoplasmic signal of our endogenous HRPA-1::GFP reporter was difficult to detect due to strong nuclear signal (Fig 4). Previously, *in vivo* proximity-based proteomics found HRPA-1 to be enriched in the nucleus hile also reporting cytoplasmic HRPA-1 localization in a few tissues, including epidermis, intestine, and body wall muscle (Reinke *et al*. 2017). Thus, it is possible that while the bulk of HRPA-1 localizes to the nucleus, HRPA-1 is present in the cytoplasm, where it may associate with the cytoplasmic miRNA machinery or the miRNA targets.

### What is the underlying mechanism for *hrpa-1* genetic interactions with miRNAs

While the physical association between HRPA-1 and miRNAs/miRISC requires future work to be resolved, we show that *hrpa-1(ok963)* genetically interacts with multiple miRNA sensitized backgrounds (Figs. 2, 3). Loss of *hrpa-1* significantly enhanced the developmental defects associated with *let-7(n2853), mir-8 mir-241(nDf51)*, and *lsy-6(ot150)* mutants (Figs. 2, 3, Table S2, S3). This suggests that HRPA-1 is normally involved in promoting miRNA activity during development and HRPA-1 activity may be broadly important for gene regulation, spanning many miRNAs’ developmental functions. HRPA-1 could affect miRNA biogenesis and/or activity and we tested the former hypothesis by assessing miRNA levels in HRPA-1-depleted animals. Surprisingly, HRPA-1 depletion did not significantly affect the levels of let-7, miR-58, or other miRNAs responsible for the developmental phenotypes observed in *hrpa-1(ok963); miRNA(reduction of function)* animals. This could indicate that HRPA-1 is not critical for biogenesis of these miRNAs but may rather be required for efficient miRNA-target repression activity (Fig 7A) or may indirectly coordinate with miRNAs to regulate developmental gene expression programs. However, it is also possible that our *hrpa-1* RNAi/IAA depletion was not sufficient to cause a wide-spread disruption in mature miRNA levels. While animals were hatched onto *hrpa-1* RNAi, the additional three-hour IAA treatment was likely not sufficient to further affect levels of the existing mature miRNAs, considering the long miRNA half-life (Miki *et al*. 2014; Vieux *et al*. 2021). Consistent with that, we observed a mild accumulation of pri-let-7 in HRPA-1-depleted RNA (1.65 fold) (Fig S5). Given the rapid processing rates of miRNA intermediates, primary miRNAs could be more responsive to dysregulation of HRPA-1-mediated processes than mature miRNAs in our approach. In addition, we did observe an enrichment for pri-let-7 in HRPA-1::FLAG IP (Fig 5E), suggesting that HRPA-1 role in primary miRNA processing could be similar to its human homolog (Guil and Ca’ceres, 2007; Michlewski and Ca’ceres, 2008). Thus, it is possible that HRPA-1 affects *let-7* processing in specific tissues and our small RNA seq was not sensitive enough to detect such tissue-specific changes in mature let-7 levels.

**Figure 7.**
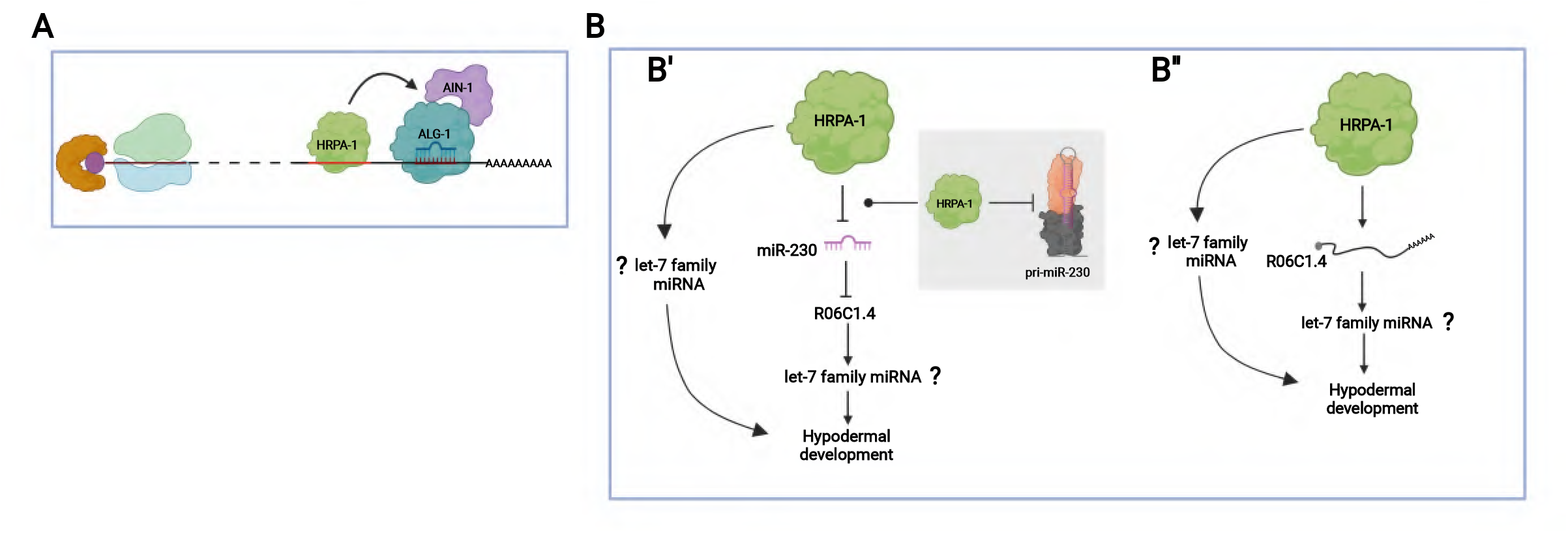
Models representing potential HRPA-1 roles in miRNA mediated gene regulation. (A) For miRNAs whose levels were unaffected by loss of *hrpa-1*, HRPA-1 could function to promote miRISC target binding and repression. (B) Models for regulation of R06C1.4 by HRPA-1 in developmental gene expression programs. (B’) Regulation of R06C1.4 could be through miR-230 processing modulation by HRPA-1. (B’’) HRPA-1 could be regulating R06C1.4 in a miRNA independent manner which could have an impact on hypodermal development via, or independent of, miRNAs. Models were drawn using BioRender (biorender.com).

### Potential gene regulatory targets of HRPA-1

Depletion of *hrpa-1* did alter levels of 20 mature miRNAs, suggesting a role in biogenesis or stability (Fig 5D), perhaps through effects on primary miRNA processing either by promoting or inhibiting microprocessor activity (Fig 7B). Interestingly, predicted targets of some of the upregulated miRNAs were downregulated in *hrpa-1* animals, supporting the hypothesis that developmental effects of *hrpa-1* loss could at least in part be brought on by miRNA biogenesis/stability misregulation.

Among the miRNA-target pairs predicted by TargetScan, depletion of HRPA-1 resulted in dysregulation of the miR-230-R06C1.4 pair (Fig 5). Mature miR-230 levels were upregulated in *hrpa-1-*depleted animals (Fig 5D), while levels of the *R06C1*.*4* mRNA went down (Fig 5H). *R06C1*.*4* knockdown partially recapitulated *hrpa-1* loss of function phenotype in miRNA mutant background (Fig 6 A,B,D). Thus, downregulation of R06C1.4 upon of depletion of HRPA-1 could be via HRPA-1 effects on miR-230 biogenesis and/or stability (Fig 7B’). HRPA-1 could also be involved in *R06C1*.*4* regulation independent of miR-230, as evidenced by enrichment for *R06C1*.*4* mRNA in HRPA-1 IP (Fig 7B’’).

Overall, we present evidence for HRPA-1 functional requirement in several miRNA-mediated developmental processes and identify potential mRNA and miRNA targets of HRPA-1, including *R06C1*.*4*. We propose HRPA-1 may interface with multiple gene regulatory machineries, similar to its human homologs. Whether *hrpa-1* coordinates with miRNAs through direct mechanisms (such as miRNA biogenesis) to regulate gene expression in *C. elegans* remains to be resolved and will be a subject of interest for future studies.

## Acknowledgements

We thank members of the Zinovyeva lab for helpful discussions and technical assistance. We are grateful to Erik Lundquist, Helge Grosshans, and Kyu Sang Joeng for sharing reagents and strains. Some of the strains used in this work were provided by the Caenorhabditis Genetics Center, which is funded by the National Institutes of Health Office of Research Infrastructure Programs P40-OD010440.

## Author contributions statement

S.H: Study concept and design, performing the experiments, analysis and interpretation of data, prepared figures, manuscript writing. G.P.P: Data analysis. A.Y.Z: Study concept and design, performing the experiments, editing manuscript, technical and material support, study supervision.

## Additional information

The authors declare no conflict of interest.

