## Supplemental materials for "hnRNPA1/2 homolog *hrpa-1* coordinates with miRNAs to regulate gene expression during *C. elegans* development"

**Table S1. List of Oligonucleotides Used in This Study**

| Oligonucleotide | Sequence |
|---|---|
| *dpy-10* crRNA | 5’-GCUACCAUAGGCACCACGAG-3’ |
| *hrpa-1* crRNA | 5’-CAGTGGGCTCATGCTCAAGG-3’ |
| *R06C1.4* crRNA1 | 5’-TAATCGATTTTTTTCTAGT-3’ |
| *R06C1.4* crRNA2 | 5’-CGCAACAGTCAAATGAACA-3’ |
| *R06C1.4::V5* donor<br>(IDT Ultramer) | 5’-<br>CCTTCAATGGACGCAACCTTCGTGTCAACTACGCCAACAAGG<br>GAGGTTCCGGTGGTTCTGGTGGATCCGGTAAGCCTATCCCAA<br>ATCCTTTGTTGGGTCTGGACTCCACGTAAACTGACAACGCCGC<br>CGAGTGACTCAAAGAGATTGCACTTG-3’ |
| *R06C1.4* genotyping primers | F: 5’-GGCCAAAACCTACTATTACC -3’<br>R: 5’-ACAAGTGCAATCTCTTTGAGTC-3’ |
| 2'-OMe miR-58 oligo | 5′: CAUCAUUGCCGUACUGAACGAUCUCAAGUC-3′ |
| 2'-OMe let-7 oligo | 5′: UCUUCACUAUACAACCUACUACCUCAACCUU-3′ |
| 2'-OMe scrambled oligo | 5′: CAUCACGUACGCGGAAUACUUCGAAAUGUC-3′ |
| qPCR pri-let-7<br>primers | F: 5’-AAGCAGGCGATTGGTGGAC-3’<br>R-5’GTTGTGAGAGCAAGACGACG-3’ |
| qPCR pri-miR-58<br>primers | F: 5’-TTACTATCCCGGCTTCAGTG-3’<br>R: 5’-CCATACCTTCCGTTCCGTAT-3’ |
| qPCR *R06C1.4*<br>primers | F: 5′-CTCCGTCTACGTTGGAAACG-3′<br>R: 5’-GTTGGCGTAGTTGACACG-3’ |

**Table S2. Summary of RNAi based phenotypic and reporter assays performed in this study.**

| miRNA mutant | *let-7(n2853)* | *mir-48 mir-241(nDf51)* | | *gfp::lin-41*[a] | | *gfp::lin-41(ΔLCS)*[b] | *cog-1::gfp*[c] | |
|---|---|---|---|---|---|---|---|---|
| | | | | wt | *let-7(n2853)* | | wt | lsy-6 |
| Phenotype / RNAi | % Animals burst through vulva (n[d]) | % Abnormal hypodermal *col-19::gfp* (n) | Seam cell # (n) | | | | | |
| **vector** | 18.3±2.2 (78) | 14.3±4 (72) | 14±0.1 (72) | 0.70±0.1 | 1.31±0.2 | 1.59 | 1.59 | 48.3±5.7 |
| ***dcr-1*** | **63.9±17.2 (122)** | **66.1±26.8 (80)** | **14.87±0.4 (80)** | 0.71±0.2 | 1.45±0.1 | 1.68 | 1.68 | n.d |
| ***hrpa-1*** | **55.6±9.7[e] (113)** | **85±7.1 (94)** | **16.7±0.5 (94)** | 0.66±0.1 | 1.23±0.1 | 1.51 | 1.51 | **62.3±3.4** |
| ***R06C1.4*** | 37.1±9.3 (86) | **44.2±4.4 (107)** | **14.5±0.5 (107)** | n.d | | n.d | n.d | |
| ***C24F3.3*** | 33.3 (52) | 6.7±1.0 (48) | 13.6±0.1 (48) | n.d | | n.d | n.d | |
| ***gcsh-1*** | 39.5 (54) | 21±4.4 (63) | 14.3±0.3 (63) | n.d | | n.d | n.d | |
| ***T10H9.5*** | n.d | 6.9±3.1 | 13.6±0.2 | n.d | | n.d | n.d | |
| ***Y57G11B.5*** | 47.7 (43) | 9.3±1.8 (68) | 13.5±0.1 (68) | n.d | | n.d | n.d | |
| ***ego-1*** | 28.2±3.2 (60) | **46.2±11.9 (55)** | **14.5±0.7 (55)** | n.d | | n.d | n.d | |
| ***ech-7*** | 24.8±8.5 (37) | 25.9±21.1 (39) | 14.2±0.9 (39) | n.d | | n.d | n.d | |

[a,b] Relative fluorescence (Green/Red) intensities for each RNAi are shown here*; n* ranges from 20 to 54

[c] Percentages of animals expressing uterine *cog-1::gfp*; *n* ranges from 88 to 110

[d] Total number of worms scored across replicates

[e] Highlighted in bold is the statistically significant value (p-value<0.05) as determined by a T-test (See Methods for details)

**Table S3. Effects of loss of *hrpa-1* and *R06C1.4* on miRNA mutant phenotypes.**

| Phenotype / Genotype | % Abnormal hypodermal *col-19::gfp* (n) | Seam cell # (n) | Alae (n) | | | % Animals with defective ASEL cell fate (n) |
|---|---|---|---|---|---|---|
| | | | Complete | gapped | no alae | |
| **wt** | 0 (62) | 12.8 (62) | 100 (45) | 0 (0) | 0 (0) | 0 (68) |
| ***mir-48 mir-241(nDf51)*** | **15 (55)**[a] | **14.0 (55)** | **50 (16)** | **37.5 (12)** | **12.5 (13)** | **n.d** |
| ***hrpa-1(ok963)*** | 0 (45) | 12.6 45) | **54 (22)** | **37.5 (15)** | **8.5 (3)** | 0 (86) |
| ***hrpa-1(ok963); mir-48 mir-241(nDf51)*** | **100 (65)** | **16.8 (65)** | **0 (0)** | **10 (4)** | **90 (34)** | n.d |
| ***R06C1.4(zen214); mir-48 mir-241(nDf51)*** | **42.3 (59)** | **14.7 (59)** | n.d | n.d | n.d | n.d |
| ***hrpa-1(ok963); lsy-6(ot150)*** | n.d[b] | n.d | n.d | n.d | n.d | **51.4 (67)** |

[a] Statistically significant values are highlighted in bold as determined using Chi-Square test.
[b] not determined

**Supplemental Figure legends**

**Figure S1. Effects of *hrpa-1* RNAi on *pdpy-30::gfp::lin-41 3'UTR* reporter expression in the vulval cells of wild type and *let-7(n2853)* L4 larvae at 20ºC.** Compromised let-7 activity in *let-7(n2853)* mutant and deletion of let-7 complementary sites (LCS) in lin-41 3' UTR leads to de-repression of *pdpy-30::gfp::lin-41 3' UTR* reporter expression levels which was unchanged upon knockdown of *hrpa-1* (quantified in B). (B) Effects of empty vector (negative control), dcr-1 RNAi (positive control), and *hrpa-1* RNAi on *pdpy-30::gfp::lin-41 3'UTR* reporter expression levels in L4 animals. Each dot represents the relative intensity in an individual worm. Vulval precursor cells used for fluorescence quantification are highlighted with dashed circles. T-test was used to determine the statistical significance.

**Figure S2. Expression levels of differentially expressed intronic miRNAs and miR-58a as well as their host genes upon HRPA-1 depletion.** Y-axis shows RPM values of miRNA expression and FPKM values of host gene expression.

**Figure S3. Functional assay in *let-7(n2853)* background using genes downregulated by depletion of HRPA-1.** Effects of RNAi knockdown of top non-collagen genes downregulated by loss of HRPA-1 on vulval bursting in wild type and *let-7(n2853)* backgrounds at 15ºC. T-test was used to determine the statistical significance.

**Figure S4. Alternative model for gene regulation by HRPA-1 and R06C1.4.** (A) R06C1.4 is a homolog of Rna15, a component of yeast cleavage and polyadenylation complex. Model for

HRPA-1 and R06C1.4 based gene regulatory mechanism in *C. elegans*. (B) Anti-FLAG based HRPA-1 IP did not capture R06C1.4::V5.

**Figure S5. Pri-let-7 level marginally increased by depletion of HRPA-1.** Fold change in pri-let-7 level as quantified by RT-qPCR in HRPA-1 depleted RNA compared to the control.

#### Figure S1

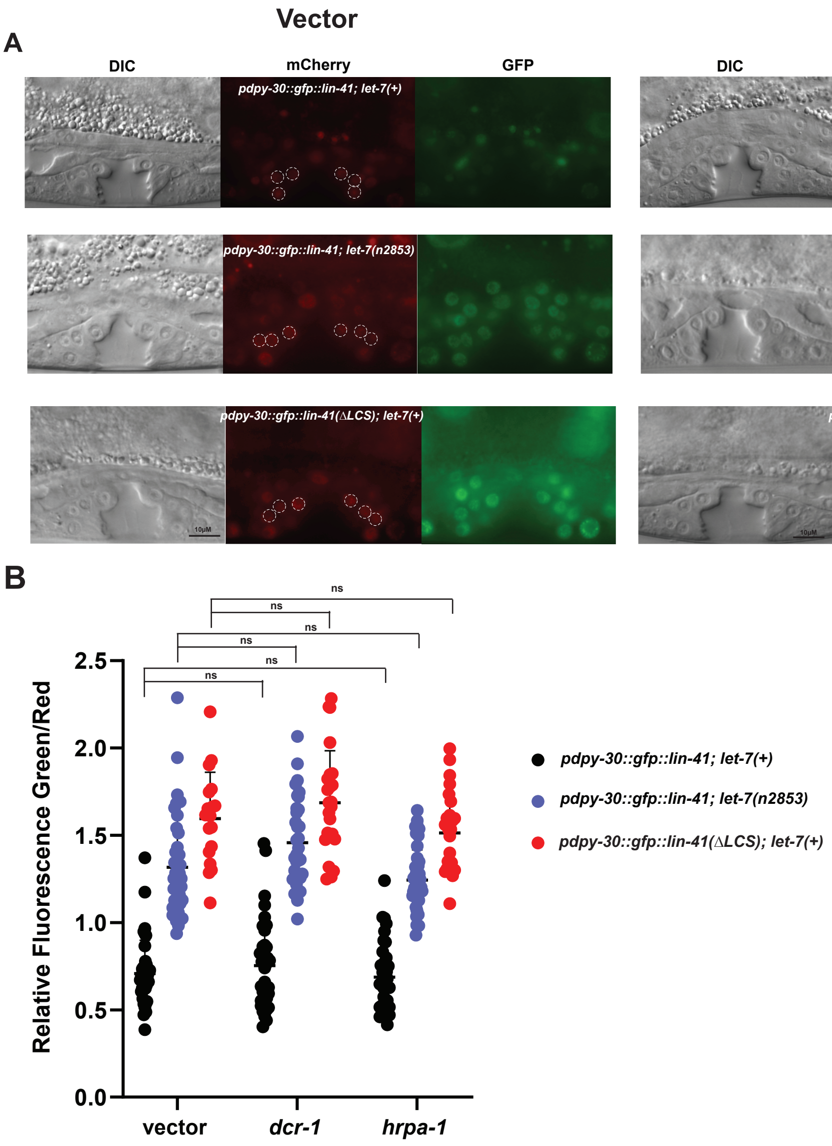

A
Vector
hrpa-1 RNAi
DIC
mCherry
GFP
pdpy-30::gfp::lin-41; let-7(+)
pdpy-30::gfp::lin-41; let-7(n2853)
pdpy-30::gfp::lin-41(ΔLCS); let-7(+)
10μM
B
Relative Fluorescence Green/Red
2.5
2.0
1.5
1.0
0.5
0.0
ns
vector
dcr-1
hrpa-1
pdpy-30::gfp::lin-41; let-7(+)
pdpy-30::gfp::lin-41; let-7(n2853)
pdpy-30::gfp::lin-41(ΔLCS); let-7(+)

#### Figure S2

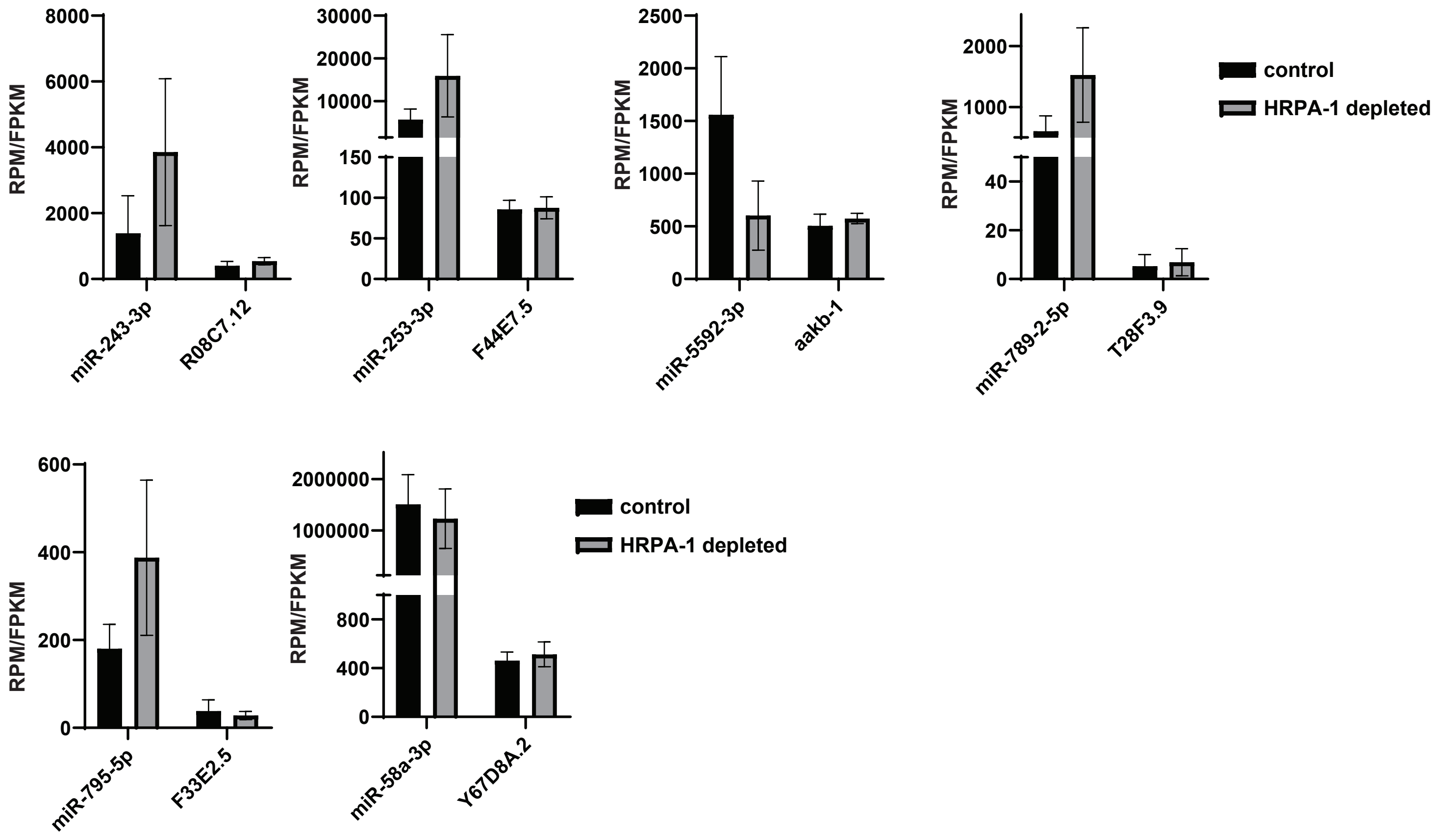
RPM/FPKM
8000
6000
4000
2000
0
miR-243-3p
R08C7.12
30000
20000
10000
150
100
50
0
miR-253-3p
F44E7.5
2500
2000
1500
1000
500
0
miR-5592-3p
aakb-1
2000
1000
40
20
0
miR-789-2-5p
T28F3.9
control
HRPA-1 depleted
600
400
200
0
miR-795-5p
F33E2.5
2000000
1000000
800
400
0
miR-58a-3p
Y67D8A.2
control
HRPA-1 depleted

#### Figure S3

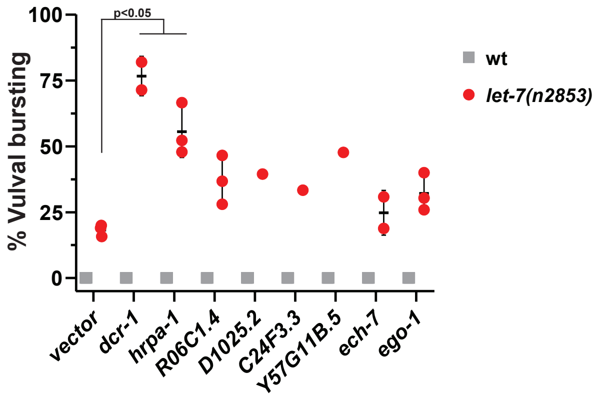

p<0.05
% Vulval bursting
100
75
50
25
0
wt
let-7(n2853)
vector
dcr-1
hrpa-1
R06C1.4
D1025.2
C24F3.3
Y57G11B.5
ech-7
ego-1

### Figure S4

**A**

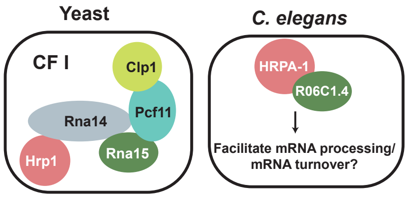

Yeast
CF I
Clp1
Rna14
Pcf11
Hrp1
Rna15
C. elegans
HRPA-1
R06C1.4
Facilitate mRNA processing/
mRNA turnover?


**B**

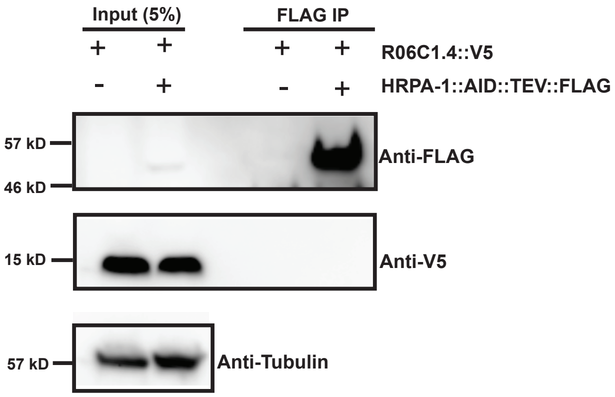

Input (5%)
FLAG IP
+ + + +
R06C1.4::V5
- + - +
HRPA-1::AID::TEV::FLAG
57 kD
46 kD
Anti-FLAG
15 kD
Anti-V5
57 kD
Anti-Tubulin

**Figure S5**

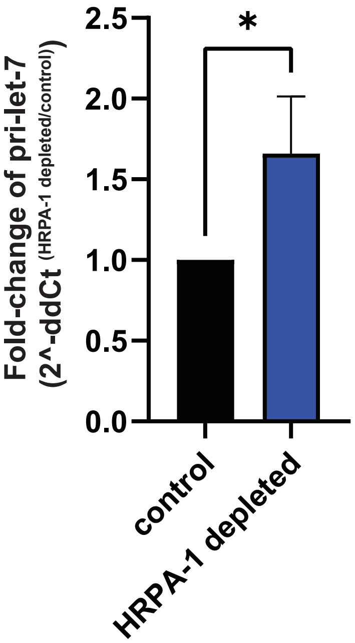

*
2.5
2.0
1.5
1.0
0.5
0.0
Fold-change of pri-let-7
(2^-ddCt (HRPA-1 depleted/control))
control
HRPA-1 depleted
